# Prion-like C-terminal domain of TDP-43 and α-Synuclein interact synergistically to generate neurotoxic hybrid fibrils

**DOI:** 10.1101/2020.12.12.422524

**Authors:** Shailendra Dhakal, Courtney E. Wyant, Hannah E. George, Sarah E. Morgan, Vijayaraghavan Rangachari

## Abstract

Aberrant aggregation and amyloid formation of tar DNA binding protein (TDP-43) and α-synuclein (αS) underlie frontotemporal dementia (FTD) and Parkinson’s disease (PD), respectively. Amyloid inclusions of TDP-43 and αS are also commonly co-observed in amyotrophic lateral sclerosis (ALS), dementia with Lewy bodies (DLB) and Alzheimer disease (AD). Emerging evidence from cellular and animal models show colocalization of the TDP-43 and αS aggregates, raising the possibility of direct interactions and co-aggregation between the two proteins. In this report, we set out to answer this question by investigating the interactions between αS and prion-like pathogenic C-terminal domain of TDP-43 (TDP-43 PrLD). PrLD is an aggregation-prone fragment generated both by alternative splicing as well as aberrant proteolytic cleavage of full length TDP-43. Our results indicate that two proteins interact in a synergistic manner to augment each others aggregation towards hybrid fibrils. While monomers, oligomers and sonicated fibrils of αS seed TDP-43 PrLD monomer aggregation, TDP-43 PrLD fibrils failed to seed αS monomers indicating selective interactions. Furthermore, αS modulates liquid droplets formed by TDP-43 PrLD and RNA to promote insoluble amyloid aggregates. Importantly, the cross-seeded hybrid aggregates show greater cytotoxicity as compared to the individual homotypic aggregates suggesting that the interactions between the two proteins have a discernable impact on cellular functions. Together, these results bring forth insights into TDP-43 PrLD – αS interactions that could help explain clinical and pathological presentations in patients with co-morbidities involving the two proteins.

## INTRODUCTION

Protein misfolding and toxic amyloid formation have come to define the pathogenesis of many neurodegenerative disorders including Alzheimer disease (AD), Parkinson’s disease (PD), amyotrophic lateral sclerosis (ALS) and frontotemporal dementia (FTD) [1, 2]. Each of these neurodegenerative pathologies is known to involve aberrant aggregation of a protein that brings to bear cellular abnormalities and toxicities. However, overlapping clinical presentations and comorbidities observed among patients with neurodegenerative diseases have motivated researchers into investigating the possibility of molecular overlaps among amyloidogenic proteins involved in them. For example, a wealth of evidence accrued over the years indicate that some of these maladies are accompanied by the deposition of common amyloid proteins such as tau and α-synuclein (αS), collectively referred to as tauopathies and synucleinopathies [3-5]. Abnormal tau inclusions are often observed alongside amyloid-β (Aβ) deposits in AD patients [6] as well as in conditions such as PD [7, 8] and prion disease [9, 10]. Similarly αS, the major protein involved in the formation of pathogenic amyloid inclusions of Lewy bodies (LBs) in PD is also observed in AD [11], dementia with Lewy bodies (DLB), and multiple system atrophy (MSA) [12, 13]. Like tau and αS, insoluble cytoplasmic inclusions of the ribonucleoprotein, tar DNA binding protein (TDP-43) are increasingly observed in many neurodegenerative pathologies such as ALS and FTD [14], as well as in patients with PD, DLB, and AD [15, 16]. There has been a surge in the reports on cross-interactions between amyloidogenic proteins such as the interactions between Aβ/tau, αS/Aβ, αS/tau [8, 17-19], and prion/Aβ [20], which have cemented the hypothesis that cross-interaction mechanisms better define the underlying cause of neurogenerative diseases and related co-pathogenesis. Along the same lines, recently it has come to light that αS and TDP-43 inclusions also co-exist in ALS, FTD and PD [15, 21]. While both αS and TDP-43 are known to independently form cytoplasmic amyloid inclusions [22-25], their co-existence raises the question of whether they interact with one another and if so, what consequence does such an interaction brings to bear. Indeed, recent reports do support this contention; TDP-43 was observed to synergistically interact and enhance αS toxicity in dopaminergic neurons [26], while immunocytochemical and immunoblot analyses on mice models and SH-SY5Y neuroblastoma cells revealed that exogenous αS fibrils promote phosphorylation and aggregation of TDP-43 [19]. Furthermore, co-expression of TDP-43 and αS was shown to induce αS pathology in *c*.*elegans* that led to significant neurodegeneration as compared to their individual expressions [27].

αS is a 140 amino acid protein containing three regions; lipid binding N-terminal domain (NTD), an aggregation-prone non-amyloid component (NAC) middle region, and an intrinsically disordered, charged C-terminal domain (CTD). NAC and a part of NTD are the primary regions responsible for aggregation, while CTD seems to play an inhibitory role for this process [28, 29]. On the other hand, TDP-43 is a 414 amino acid containing ribonucloprotein protein with an N-terminal domain, two RNA recognition motifs (RRMs), and a disordered prion-like C-terminal domain (PrLD) containing low complexity sequences [30]. TDP-43 is known to be involved in transcriptional regulation and RNA splicing [31, 32]. In pathology, the protein is translocated into the cytoplasm where it undergoes aberrant proteolytic cleavage that generates different C-terminal fragments (CTFs; C35, C25, and C18), which consequently form toxic insoluble inclusions [33-38]. Furthermore, under stress conditions, cytoplasmic TDP-43 undergoes liquid-liquid phase separation (LLPS) by coacervating with RNA and other proteins to form membraneless stress granules (SGs) [39, 40]. If and how SGs play a role in the formation of cytoplasmic TDP-43 inclusions and cellular toxicity remains unclear. PrLD, a segment of such pathological aggregates, is primarily known to drive the fibrillization process and mediates protein-protein interactions [41, 42]. Despite multiple factors in regulating fibrillization of TDP-43 PrLD, a common biophysical phenomenon, liquid-liquid phase separation (LLPS) is also recently known to modulate the aggregation process via electrostatic interactions [40].

Here, we sought to determine how αS and TDP-43 interact with one another to exacerbate cellular toxicity, by investigating the full-length αS and TDP-43 PrLD as PrLD forms a key part of pathogenic proteolytic products of TDP-43 (Figures 1a and 1b). Our results indicate that αS and TDP-43 PrLD monomers, oligomers or fibrils cross-interact and modulate their aggregation behavior. Moreover, αS also enhance TDP-43 PrLD aggregation by modulating the liquid droplets formed by the phase separation of TDP-43 PrLD in presence of RNA. In all, cross-seeded aggregates show higher cytotoxicity than the individual aggregates suggesting the potential for such mechanisms to present greater pathogenicity.

**Figure 1.**
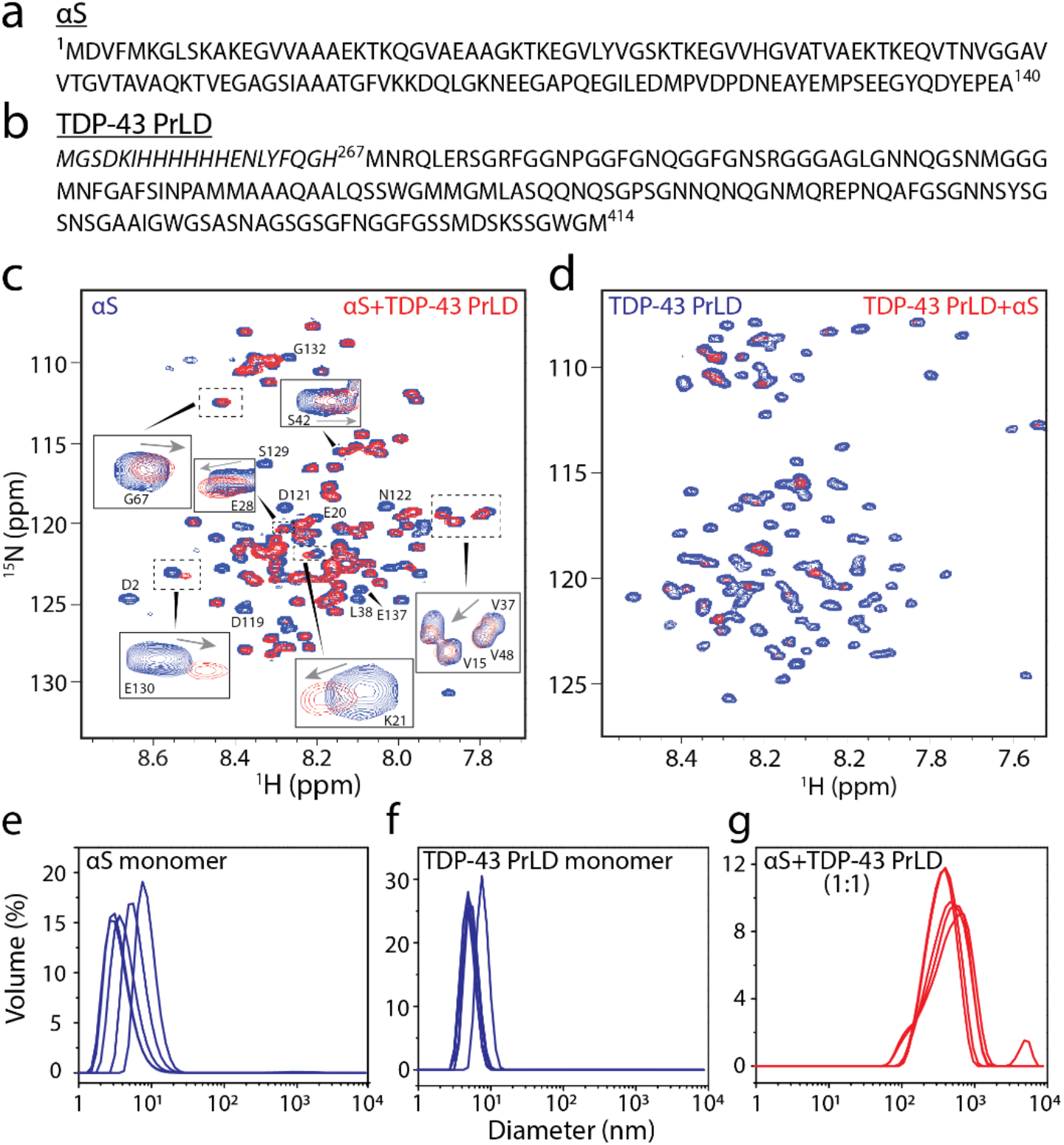
^1^H-^15^N HMQC spectroscopy of αS and TDP-43 PrLD along with DLS of respective reactions. a-b) Sequence of full-length 1-140 amino acid residues αS (a) and 267-414 amino acid residues TDP-43 PrLD used in the experiment. c) ^1^H-^15^N HMQC of 10 μM ^15^N-labeled αS monomers alone (blue) or in the presence of unlabeled 10 μM TDP-43 PrLD monomers (red) in 20 mM MES buffer pH 6.0 at 37 °C. d) ^1^H-^15^N HMQC of 10 μM ^15^N-labeled TDP-43 PrLD monomers alone (blue) or in presence of unlabeled αS (red). e-g) DLS histograms of 10 μM αS monomer control, TDP-43 PrLD monomer control and co-incubated mixture 1:1 αS and TDP-43 PrLD in the same buffer and temperature conditions taken within 10 minutes of incubation.

## RESULTS

### αS and TDP-43 PrLD monomers synergistically promote fibrillizatio

First, we questioned whether monomers of αS and TDP-43 PrLD are able to interact with one another. To answer this, heteronuclear multiple quantum coherence (HMQC) spectroscopy was performed on a uniformly ^15^N labeled αS and TDP-43 PrLD individually. Both proteins showed narrow chemical shift dispersions in the amide region (7.8-8.7 ppm for αS and 7.7-8.5 for TDP-43 PrLD in the ^1^H dimension) confirming the well-known disordered structures for both the proteins (blue; Figures 1c and 1d). To see whether the two proteins interact with one another, HMQC spectra of the ^15^N-enriched proteins were observed upon incubating with unlabeled, natural isotope-abundant proteins in equimolar concentrations at 37 °C. The spectrum of αS co-incubated with unlabeled TDP-43 PrLD showed the disappearance of several cross-peaks along with significant changes in the chemical shifts (Figure 1c). The disappearance of chemical shifts is largely attributed to the aggregation of proteins which leads to significant line broadening and/or loss of signal intensities [43, 44]. Interestingly, shifts in cross-peaks were largely confined to the N-terminal and C-terminal ends of αS (residues 1-40 and 100-40; boxes, Figure 1c) while the disappearance of cross-peaks corresponded to the central amylodogenic NAC of the αS (residues ∼61-95; Figure 1c). Considering that the N- and C-terminal domains are charged and NAC domain of αS is prone to amyloid formation, it is likely that TDP-43 PrLD induces aggregation of the NAC domain αS. Similarly, a substantial signal loss of cross-peaks was observed for TDP-43 PrLD upon co-incubation with αS indicating potential aggregation of the former (Figure 1d). In order to confirm the NMR observations, aggregation of the co-incubated samples was analyzed by dynamic light scattering (DLS) analysis. The control monomers of αS and TDP-43 PrLD showed a hydrodynamic diameter of ∼ 10 nm individually (Figure 1e and f). In contrast, the co-incubated samples showed aggregation with an average diameter of >10^3^ nm within 10 minutes of incubation (Figure 1g). These results confirm the NMR observations that αS and TDP-43 PrLD interact with one another to promote high molecular weight aggregates.

To further understand the mechanism of αS and TDP-43 PrLD interactions, we set forth to investigate the aggregation kinetics using thioflavin T (ThT) fluorescence assay (Figure 2). In one reaction set, αS concentration was held constant at 20 μM while TDP-43 PrLD concentrations were increased from 1, 5, 10 and 20 μM (0.05, 0.25, 0.5 and 1.0 molar equivalents). The reactions were incubated at 37 °C and the kinetics of aggregation was monitored for 48 hours (Figure 2a). The control αS did not show any aggregation during this period (△; Figure 2a). While TDP-43 PrLD alone in the absence of αS showed concentration-dependent aggregation (○; Figure 2a), the co-incubated samples with αS showed decreased lag times and greater ThT intensities (● Figure 2a) suggesting synergistic augmentation of aggregation between the two proteins. Alternatively, to investigate whether αS can modulate the aggregation behavior, TDP-43 PrLD was held constant at 20 μM and αS concentrations were varied from 0.05-, 0.5-, 1- or 2-fold molar equivalence corresponding to 1, 10, 20 or 40 μM, respectively (Figure 2b). TDP-43 PrLD control aggregated with lag time of 8 h (○; Figure 2b) while αS did not show aggregation in the 50 h (△; Figure 2b). concentrations. However, the addition of increasing molar ratio of αS resulted in decreased TDP-43 PrLD lag time of aggregation but with a proportional increase in ThT fluorescence intensity (● Figure 2b).

**Figure 2.**
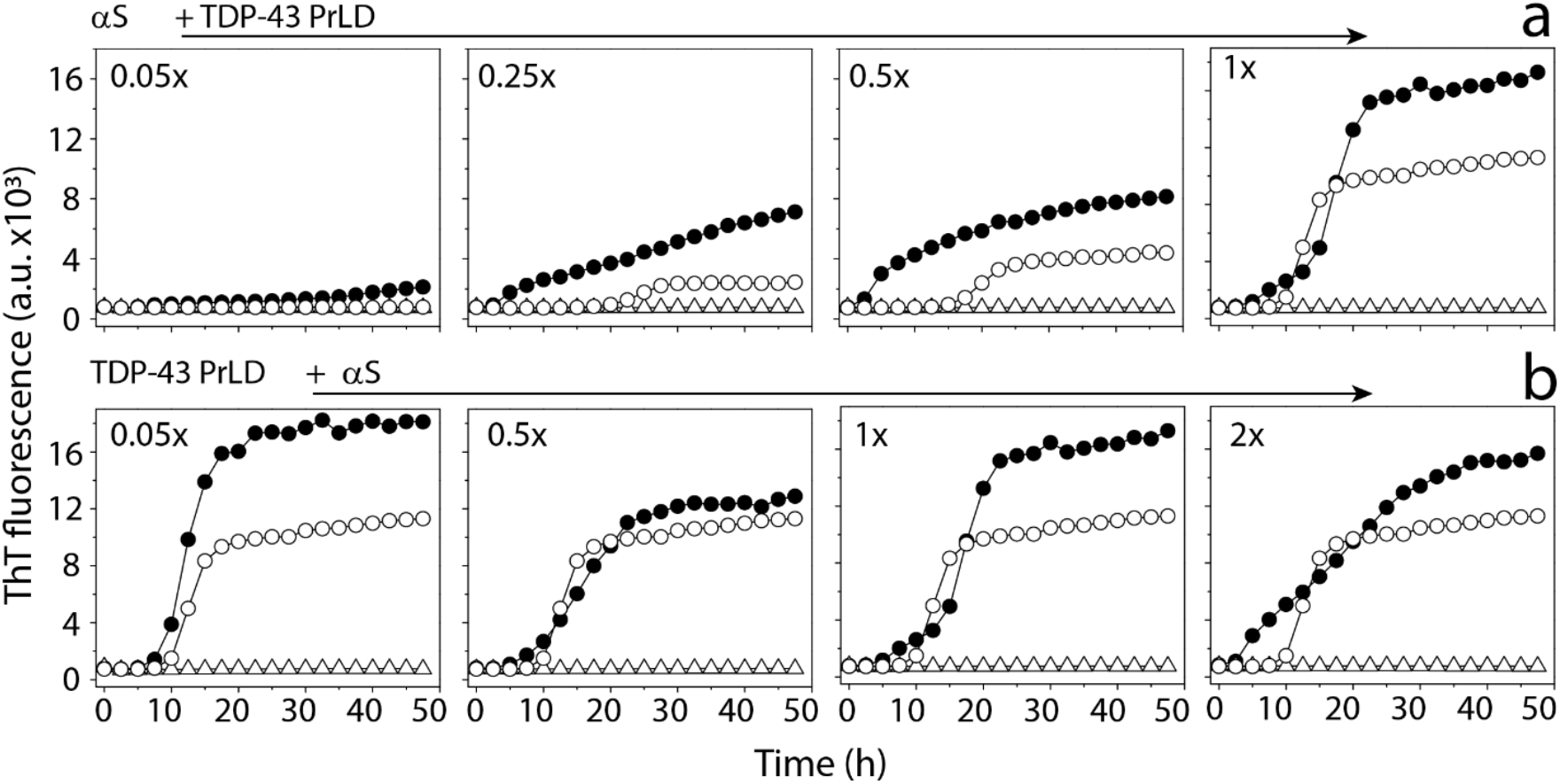
ThT aggregation kinetics of monomeric αS and TDP-43 PrLD in 20 mM MES buffer at pH 6.0 a) ThT fluorescence of 20 μM αS △in presence of 0.05 to 1 molar ratio of TDP-43 PrLD to αS ● and respective TDP-43 PrLD controls (○). b) ThT fluorescence of 20 μM TDP-43 PrLD (○) in presence of 0.05 to 2 molar ratio of αS to TDP-43 PrLD ● and respective αS controls △.

In order to monitor the effect of sub-stochiometric incubations of one protein on another, higher concentration of αS at 50 μM was incubated in presence of 5 μM (0.1x molar equivalents) and 2 μM (0.04x molar equivalents) TDP-43 PrLD. The reverse incubations with a higher concentration of TDP-43 PrLD were prohibitively difficult due to the increased aggregation propensity of the protein that showed lag time of smaller than three hours for 50 μM TDP-43 PrLD (data not shown). The reactions containing co-incubations of 50 μM αS with sub-stoichiometric TDP-43 PrLD showed significant decreases in lag time of αS fibrillization as compared to either αS or TDP-43 PrLD controls (Figure 3a). The co-incubation with 0.1 and 0.04 molar equivalences showed lag times of 10 and 20 hours, respectively (Figure 3a) suggesting TDP-43 PrLD promotes significant aggregation of αS. After 72 hours of incubation, reaction samples were centrifuged at 18,000xg for 20 minutes to sediment fibrils formed, if any. Both the samples before centrifugation and supernatant of centrifuged samples were then subjected to immunoblot analysis using monoclonal αS antibody (Figure 3b). The blot showed the presence of a band corresponding to > 260 kDa that failed to enter the gel in both co-incubated samples (T; Figure 3b) in addition to monomeric, dimeric, and trimeric αS that was present in all the samples. The corresponding supernatant samples did not contain this high molecular weight band (S; Figure 3b) which confirms that the high molecular weight species are sedimentable aggregates or fibrils. To further confirm the presence of αS fibrils in the co-incubated samples, differential interference contrast (DIC) microscopy analysis of the fibrils was employed. The pellet obtained from the samples centrifuged after 72 hours was resuspended in 20 mM MES buffer pH 6.0 and incubated with 10 μM thioflavin-S (ThS) and imaged after 10 minutes. The images revealed the presence of higher ThS positive fibrils in the reaction compared to TDP-43 PrLD and αS controls (Figure 3c). Further quantitative analysis was carried out to assess the amount of proteins present in the fibrils using MALDI-ToF. Relative quantitation was performed using a known amount of cytochrome C (1.42 μM) as an external standard that was co-analyzed with the samples in the mass spectrometer (Figure S1). Analysis of the data obtained revealed that the relative amounts of αS in the pellet of reactions containing 0.1 and 0.04 molar equivalence of TDP-43 PrLD were higher compared to the individual protein controls (Figure 3d). However, αS amount in the supernatant of the control reaction was significantly higher compared to the reactions (data not shown), indicating proportional increase in αS fibrillization on co-incubation with TDP-43 PrLD. Similar relative quantification of TDP-43 PrLD to standard in the pellets of reaction showed a higher amount of TDP-43 PrLD than respective control reactions indicating enhanced fibrillization in presence of αS (Figure 3e), although the absolute amount of TDP-43 PrLD in this samples was lower to αS attributable to the lower concentrations in the incubations. The data also indicate that the sedimented fibrils contain both αS and TDP-43 PrLD suggesting hybrid fibrils. Taken together, the data indicate that both αS and TDP-43 PrLD monomers interact with one another and promote fibrillization synergistically.

**Figure 3.**
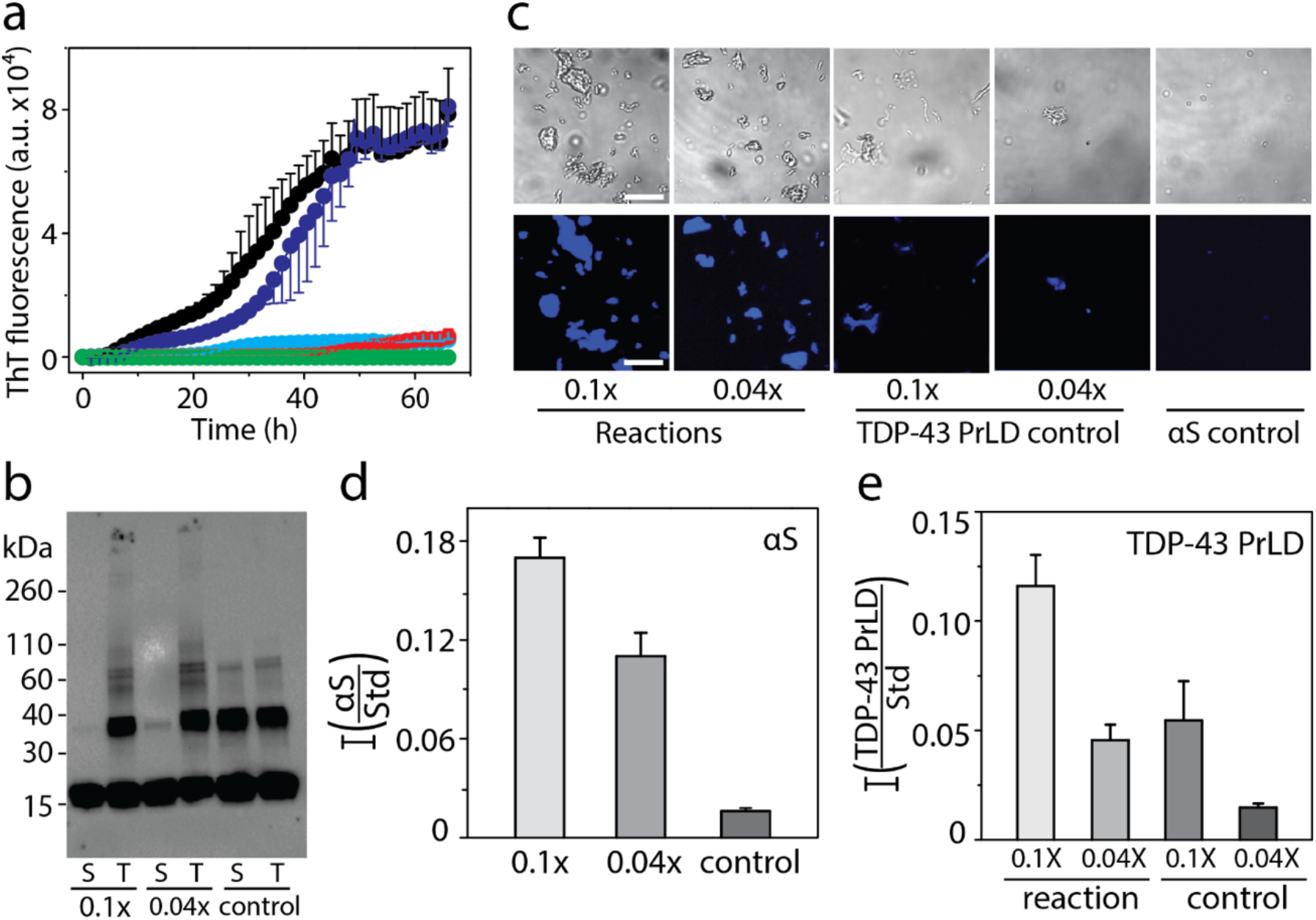
Interaction of monomeric TDP-43 PrLD and αS in 20 mM MES buffer pH 6.0. a) ThT fluorescence kinetics of 50 μM αS alone 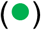 and in presence of 0.04 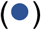 and 0.1 (●) molar ratio of TDP-43 PrLD to αS, and respective 0.04 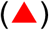 and 0.1 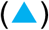 TDP-43 PrLD controls. b) Western blot of ThT reactions using αS antibodies. Aliquot of sample from the reaction at 72 hours was subjected to western blot as total sample (T), and supernatant (S) after centrifuging at 18,000 xg c) Representative fluorescence microscopic images of Thioflavin S (ThS) stained αS and TDP-43 PrLD aggregation reactions at 72 hours after centrifuging at 18,000 x g (Scale bar = 20 μm). d-e) Relative quantification of αS and TDP-43 PrLD to the Cytochrome C internal standard in pellet of reactions and control at 72 hours after centrifugation at 18,000 xg; 0.1x and 0.04x indicates molar ratio of TDP-43 PrLD to αS.

### αS oligomers and sonicated fibrils seed fibrillation of TDP-43 PrLD monomers but αS monomers are innocuous to seeding by TDP-43 PrLD sonicated fibrils

Having established that monomers of αS and TDP-43 PrLD interact and promote each other’s aggregation, we questioned if their interactions are restricted to only monomers or whether oligomers and sonicated fibrils can also cross-seed respective monomers. To answer these questions, dihydroxyphenyl acetaldehyde (DOPAL), a metabolite of dopamine biogenesis, induced oligomers of αS were used as *de facto* oligomers along with sonicated fibrils of both αS and TDP-43 PrLD. Isolation of stable oligomers of TDP-43 PrLD was not successful and hence were excluded from the investigation. DOPAL-derived oligomers of αS were generated and isolated by size exclusion chromatography (see Materials and Methods). The αS oligomers showed a monodisperse peak in dynamic light scattering (DLS) around 10 nm diameter (Figure 4a) with a molecular weight centered at ∼ 48 kDa (∼ 3mer) observed in immunoblot along with two faint bands corresponding to ∼ 33 and 64 kDa species (2 and 4mers, respectively) (inset; Figure 4a). Moreover, circular dichroism (CD) spectra of the αS oligomers showed random coiled structure with characteristic minima at 195 nm (Figure 4b) while morphologically they are seen as punctate spheres of ∼4-6 nm height by atomic force microscopy (AFM) (inset; Figure 4b). Incubation of these oligomers at 0.5 and 1 μM concentrations with 15 μM of monomeric TDP-43 PrLD in 20 mM MES at pH 6.0 resulted in decrease in lag time of TDP-43 PrLD aggregation to ∼ 6h (Figure 4c). Immunoblot analysis of the samples after 12 h showed the formation high molecular weight fibril bands that is absent in the control sample of TDP-43 PrLD (inset; Figure 4c), clearly indicating that αS oligomers are able to seed TDP-43 PrLD monomers. Seeding of 20 μM monomers of either αS or TDP-43 PrLD with 1, 2 or 4 μM sonicated fibrils of TDP-43 PrLD or αS, respectively showed different behavior. Homotypic seeding of αS monomers by αS sonicated fibrils showed an immediate increase in ThT intensities as expected for seeding by elongation mechanism [45] (Figure 4d). In contrast, the seeding of TDP-43 PrLD monomers by αS sonicated fibrils displayed a more sigmoidal response (Figure 4d), but with a shorter lag time of ThT fluorescence to that of the TDP-43 PrLD control. This suggests that TDP-43 PrLD monomers may undergo conformational alterations before they grow on αS fibrils. Similarly, homotypic seeding of TDP-43 PrLD monomers by TDP-43 sonicated fibrils showed immediate and substantial increases in ThT intensities (Figure 4e). But in stark contrast, seeding of αS monomers by TDP-43 PrLD sonicated fibrils failed to show interactions (Figure 4e). The secondary structure of the cross-seeded reactions from (c-e) was analyzed by FTIR (Figure 4f). The cross-seeded reaction of TDP-43 PrLD monomers with αS sonicated fibrils or DOPAL-derived αS oligomers displayed a parallel β-sheet structure with an absorbance peak centered at 1615 cm^-1^ (pink and black; Figure 4f) as opposed to the monomeric samples that show predominantly random coil structure at 1643 cm^-1^ or 1650 cm^-1^ (blue and red; Figure 4f). In contrast, seeding of αS monomers with TDP-43 PrLD fibrils showed a predominant random coil structure at 1643 cm^-1^ (green; Figure 4f) suggesting minimal or no seeding. Finally, morphological features of the seeded aggregates were investigated by AFM. Sonicated fibrils of αS and TDP-43 PrLD showed fragmented aggregates as expected (Figures 4g and 4i). TDP-43 PrLD monomers seeded with sonicated αS fibrils after 8h of incubation showed smooth fibrils (Figure 4h). However, αS monomers seeded with TDP-43 PrLD fibrils did not show fibers after 8h of incubation (Figure 4j). These results are in agreement with kinetics data supporting the observation of the difference in the ability to seed. Together, these data suggest that while TDP-43 PrLD monomers seem to be amenable for seeding by different αS species, αS monomers are selective towards TDP-43 PrLD monomers and not sonicated fibrils.

**Figure 4.**
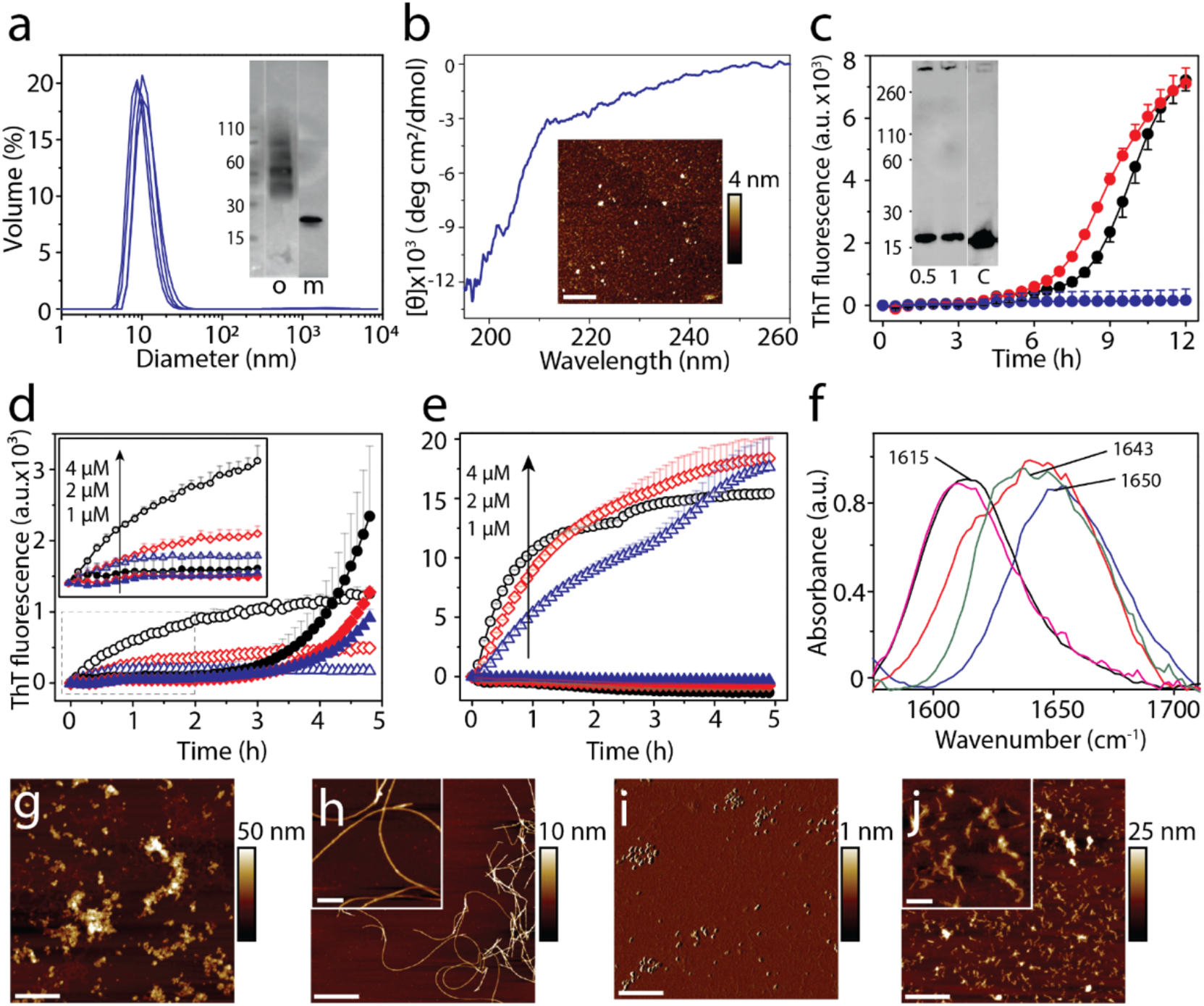
Cross-seeding of oligomers and sonicated fibrils with monomers. a) DLS analysis of DOPAL-derived αS oligomers isolated from SEC. (inset) SEC fraction containing the oligomer ‘o’ used in the study; ‘m’ refers to control monomer. b) CD spectra and AFM height image (inset) of DOPAL-derived αS oligomers used for cross-seeding reaction (scale bar = 1μm). Cross-seeding reactions with monomers of 15 μM TDP-43 PrLD alone 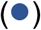 or in the presence of 0.5 μM (●) or 1 μM 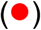 DOPAL-derived αS oligomers in the presence of 10 μM ThT. (inset) immunoblot of the reaction after 12 h probed with TDP-43 antibody; ‘c’ refers to TDP-43 PrLD monomer control. d) Seeding of 20 μM αS monomers with 4 μM (○), 2 μM 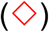, and 1 μM 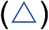 of αS sonicated fibrils, and 20 μM TDP-43 PrLD monomers seeded with 4 μM (●), 2 μM 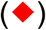, 1 μM 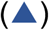 of αS sonicated fibrils. e) Seeding of 20 μM TDP-43 PrLD monomers with 4 μM (○), 2 μM 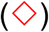, and 1 μM 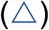 of TDP-43 PrLD sonicated fibrils and seeding of 20 μM αS monomers with 4 μM (●), 2 μM 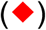, 1 μM 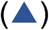 of TDP-43 PrLD seed. f) FTIR analysis of cross-seeding reactions from (c-e). TDP-43 PrLD monomers cross-seeded with 1 μM DOPAL-derived αS oligomers after 12 h of incubation (—); TDP-43 PrLD monomers seeded with 1 μM αS sonicated fibrils 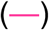; αS monomers seeded with 1 μM TDP-43 PrLD sonicated fibrils (—) along with controls such as αS monomers 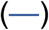 and TDP-43 PrLD monomers 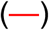 (g-j) AFM height image of αS sonicated fibrils (g), αS sonicated fibrils seeded TDP-43 PrLD monomers (h), TDP-43 PrLD sonicated fibrils (i), and TDP-43 PrLD sonicated fibrils seeded αS monomers (j) (scale bar = 1 μm, inset = 200 nm).

### αS modulates LLPS of TDP-43 PrLD and RNA and induces insoluble aggregates

One of the pivotal roles of TDP-43 in pathophysiology is that under stress conditions they are known to coacervate with RNA and other proteins to undergo LLPS to form stress granules (SGs) in the cytoplasm [39, 46]. To see whether αS is able to modulate LLPS, TDP-43 PrLD was labeled with Hilyte-647 and mixed with unlabeled TDP-43 PrLD at 1% molar ratio to a final total concentration of 20 μM protein. Similarly, αS was labeled with Hilyte-405 and mixed with unlabeled protein in 1:99 ratio, and this sample was used at final concentrations of 5 and 1 μM in 20 mM MES buffer at pH 6.0 at 37 °C. As expected, TDP-43 PrLD spontaneously phase separates in presence of RNA to form liquid droplets (0h; Figure 5a). The addition of αS led to instant changes in the morphology of the droplets to become somewhat distorted, non-spherical structures in both stoichiometric incubations (0h; Figure 5a). The same reactions monitored after 24 hours showed the droplets becoming more distorted and aggregated as opposed to the control which continued to have well-defined phase-separated liquid droplets (24 h; Figure 5a). To investigate whether the droplet deformity is due to aggregation, the dynamics of liquid droplets were probed by FRAP analysis. The control TDP-43 PrLD – RNA droplets showed significant FRAP recovery rates both at 0 h and 24 h indicating that the samples retained significant fluid characteristics (Figure 5b and c). However, the FRAP recovery rates of αS co-incubated TDP-43 PrLD droplets reduced significantly at 0 h which remained constant over the period of 24 h with a more dramatic effect with 5 μM than 1 μM αS (Figure 5b and c), which suggest gelation or aggregation samples. To unambiguously confirm if the attenuation of droplet fluidity is due to aggregation, the same reactions were monitored by ThT. The data indicated that the ThT fluorescence did not show much increase in 24 h for the TDP-43 PrLD–RNA control reaction (○; Figure 5d). In contrast, samples co-incubated with αS showed increased ThT fluorescence suggesting aggregation of TDP-43 PrLD as observed in other reactions (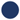 and 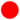; Figure 5d). Taken together, these data suggest that αS modulate phase separation of TDP-43 PrLD with RNA and enhance its aggregation.

**Figure 5.**
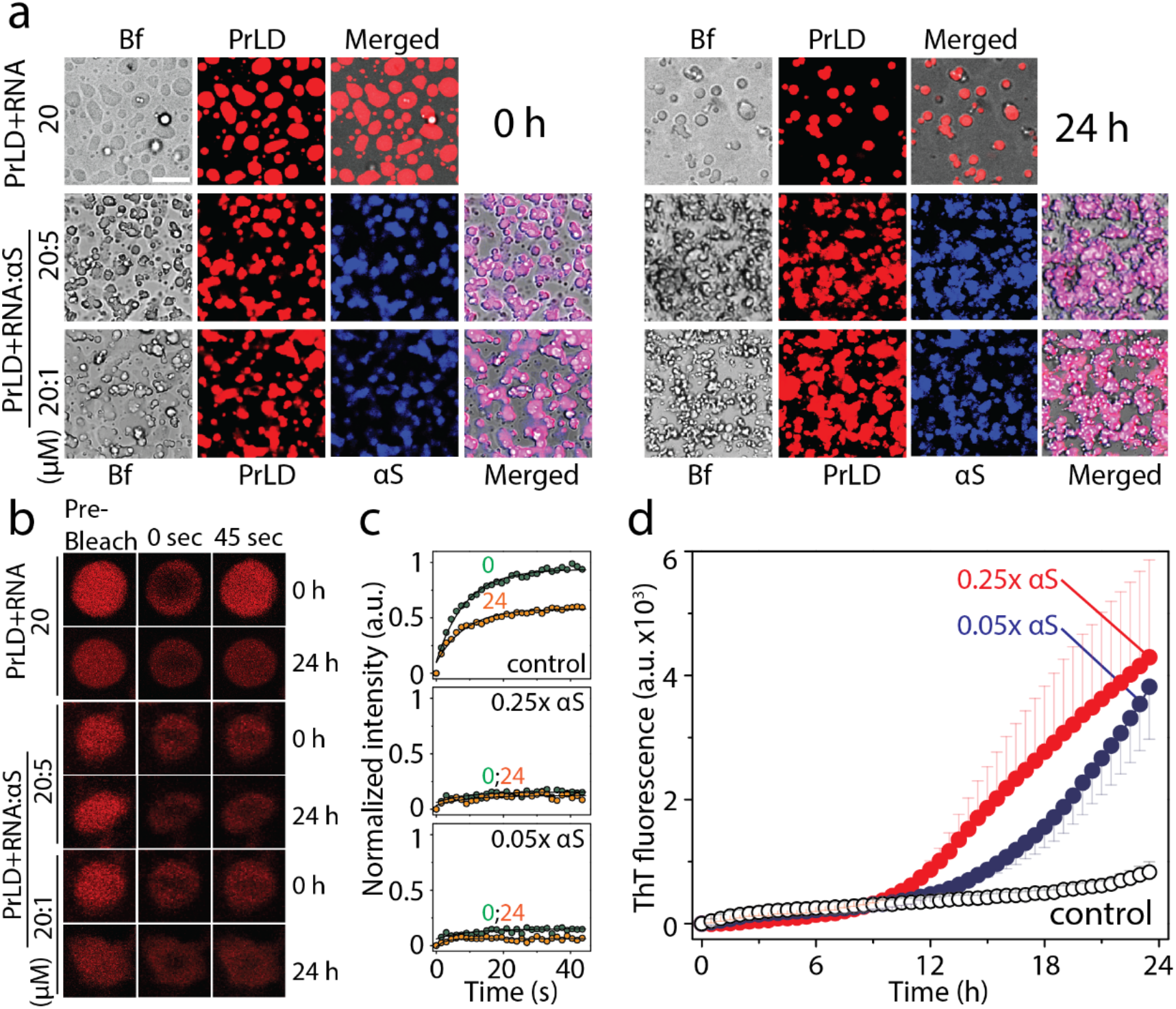
Modulation of TDP-43 PrLD LLPS by αS. a) Timestamped confocal images of the co-incubations of Hilyte-647 labelled TDP-43 PrLD (20 μM) and RNA (40 μug/mL) in the absence and presence of Hilyte 405 labelled αS (5 or 1 μM) in 20 mM MES buffer pH 6.0 at 37 °C; Bf represents ‘bright field’. b) FRAP analysis on the selected droplets from the reactions before (pre-bleach), during (0 sec), and after photobleaching (45 sec), immediately after incubation (0 h) and after 24 h. c) Normalized kinetics of fluorescence recovery data obtained from FRAP intensity; TDP-43 PrLD and RNA control reaction along with 5 μM and 1 μM αS at 0h 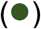 and after 24 h 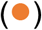. The data was fit to a first order exponential growth equation (solid lines). d) Corresponding ThT fluorescence kinetics of the reactions; TDP-43 PrLD and RNA (○) control reaction along with sub-stoichiometric, 1 μM 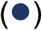 or 5 μM 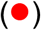 αS incubations.

### Cross-seeded heterotypic fibrils are more cytotoxic than the homotypic fibrils of αS or TDP-43 PrLD

Recently, it was found that TDP-43 enhances the toxicity of αS in dopaminergic neurons resulting in neurodegeneration [26]. Thus, we wanted to investigate and compare the toxicities of heterotypic fibrils formed by cross-seeding and the homotypic ones formed by individual αS or TDP-43 PrLD fibrils in human neuroblastoma SH-SY5Y cells. In order to accomplish this, 20 μM αS monomers were co-incubated with 5 μM TDP-43 PrLD monomers (TDP-43 PrLD: αS = 1:4) in quiescent conditions for 24 hours at 37 °C. Similarly, 20 μM TDP-43 PrLD monomers were co-incubated with 5 μM αS monomers (TDP-43 PrLD: αS = 4:1) under identical conditions. The samples were then diluted five-folds and incubated with freshly cultured SH-SY5Y cells for additional 24 hours. Appropriate control reactions corresponding to homotypic proteins were also set up and incubated under similar conditions. Cell viability of the confluent cells was then determined by XTT assay (see Materials and Methods) [47]. The results indicate that in both stoichiometries, only the co-incubated samples showed higher toxicity than the homotypic aggregates (Figure 6). Among the two, 1:4 reaction of TDP-43 PrLD: αS monomers showed a substantially greater degree of toxicity (∼40%) as compared to the 4:1 sample (∼28%), suggesting that the effect of sub-stoichiometric addition of TDP-43 PrLD enhances αS aggregation and toxicity. In case of cross-seeding of fibrils to monomers, αS fibrils seeded TDP-43 PrLD aggregation showed a substantial increase in toxicity (∼41%) compared to the αS fibril seeds or TDP-43 PrLD monomers (< 20%) (Figure 6). TDP-43 PrLD fibril seeded αS aggregation did not show an increase in toxicity compared to the controls. These results correlate with the kinetics and morphology of the seeding reaction in which only αS fibrils seeded TDP-43 PrLD aggregation showed aggregation (Figure 4). Similarly, the addition of αS monomers to the liquid droplets of TDP-43 PrLD and RNA that promoted insoluble fibrils also showed an increase in toxicity (∼30%) as compared to TDP-43 PrLD-RNA control (< 20%). In contrast to these results, only αS oligomer (DOPAL-induced) seeded TDP-43 PrLD fibrils did not show a statistically significant increase in toxicity (Figure 6). One of the main reasons for this result is due to the fact that αS oligomer is DOPAL-induced and not a bonafide aggregation pathway intermediate; clearly, DOPAL-induced αS oligomers showed maximum toxicity (∼57%; Figure 6). Therefore, it is possible that fibrils of TDP-43 PrLD seeded by these oligomers do not display any higher degree of toxicity. Together, these data suggest that the synergistic interactions between TDP-43 PrLD and αS lead to the formation hybrid fibrils which are more toxic than those of their individual proteins.

**Figure 6.**
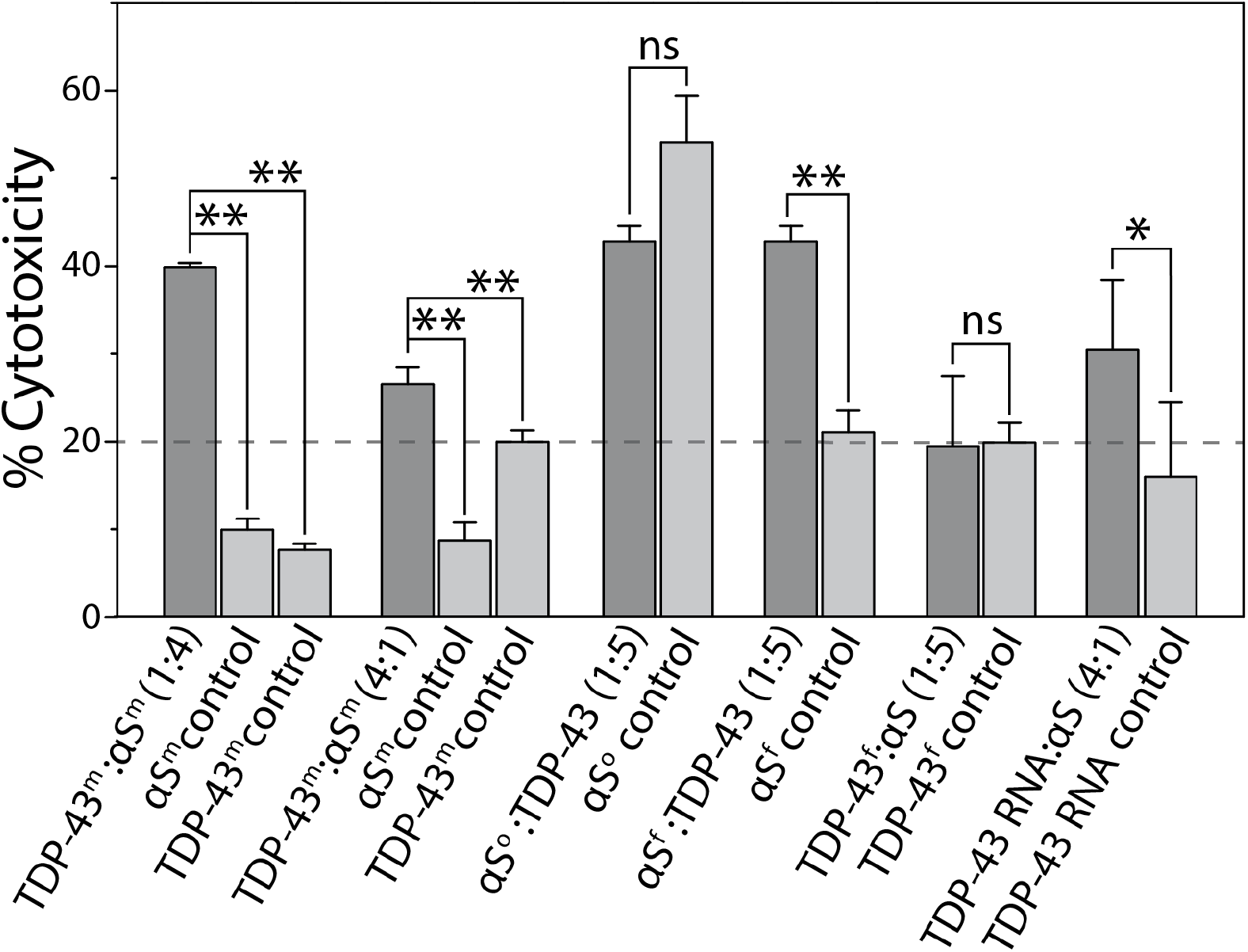
Cytotoxicity of monomeric αS and TDP-43 PrLD species along with homotypic and heterotypic fibrillar species of αS and TDP-43 in SH-SY5Y cells by XTT assay. ‘m’, ‘o’ and ‘f’ in the superscript represents monomer, oligomer, and sonicated fibril respectively. All the data were obtained in triplicates, * represents p<0.1 and **represents p<0.01 based on one-way ANOVA analysis.

## DISCUSSION

The work reported here is focused on the effects of TDP-43 PrLD on αS and vice versa. We chose to focus on PrLD instead of the full-length TDP-43 for a variety of reasons; First, the post-mortem ALS and FTD brain tissues show an abundance of TDP-43 CTFs ranging between 35 and 18 kDa (35, 25, 20, and 18 kDa) [48-51], all of which constitute PrLD (17 kDa) a major part. In a more recent study on mass spectrometric analysis of post-mortem brain samples, enhanced levels TDP-43 C-terminal truncation fragment of 266-414 (corresponds to TDP-43 PrLD) was observed in ALS and AD patients [38]. Second, TDP-43 CTFs are primarily involved in the formation of cytoplasmic inclusions generated upon aberrant proteolytic cleavage of the full-length TDP-43 under pathological conditions in which PrLD plays a major role [33, 42, 52]. Third, CTFs ranging between 35 and 18 kDa are also produced by alternative splicing of TDP-43 mRNA and are linked to pathology [51, 53]. Fourth, the PrLD of TDP-43 is known to play a major role in SG formation, aggregation as well as protein-protein interactions [54-56]. Lastly, almost all pathogenic mutations lie within PrLD [42, 57] implying its significance in pathophysiology.

The results presented here demonstrate synergistic interactions between TDP-43 PrLD and αS, which in part, recapitulate the pathobiological observations. The results also bring out mechanistic differences in their interactions; the data indicate that the three key αS species such as monomers, oligomers and sonicated fibrils are indiscriminate in interacting with and getting modulated by TDP-43 PrLD monomers (Figure 7). However, only monomeric TDP-43 PrLD, and not the sonicated fibrils, seem to preferentially interact with αS monomers and aggregates (Figure 7). From these results, one may conjecture that early stages of TDP-43 proteinopathies may be susceptible to modulation by the presence of αS aggregates in the form of Lewy bodies. Furthermore, the interaction of TDP-43 PrLD and αS monomers seem to be cooperative with both assisting one another in promoting high molecular weight, hybrid aggregates expeditiously. The hybrid aggregate presence was confirmed by the presence of both proteins in the sedimented fibrils of the co-incubated samples (Figure S2). The effect of TDP-43 PrLD on αS is far more pronounced than vice versa. This is partly due to αS being relatively slow to aggregate as compared to TDP-43 PrLD; for example, the lag times for αS and TDP-PrLD (20 μM) aggregation at 37 °C and in identical quiescent buffer conditions is, > 7 days and 8 hours, respectively (data not shown). Therefore, augmentation of αS aggregation by TDP-43 PrLD is readily comprehensible as the lag time of aggregation is reduced to 20 hours from weeks (Figure 3a). On the other hand, the effect of αS on TDP43 PrLD is more subtle yet discernable (Figure 1b). This is especially true with DOPAL-derived αS aggregates which augment TDP-43 PrLD aggregation significantly (Figure 4c). αS sonicated fibrils seeding of TDP-43 PrLD monomers show a cooperative mechanism with a delay in elongation, which could indicate conformational reorganization of TDP-43 monomers induced by αS sonicated fibrils. On the other hand, TDP-43 PrLD sonicated fibrils failed to interact with αS monomers suggesting possible incompatibility of TDP-43 PrLD sonicated fibrils structure to seed. Clues about the structural basis of interactions come from the NMR data, which shows that the TDP-43 PrLD seem to preferentially interact with αS on the N-terminal and C-terminal ends of the protein (1-40 and 100-140, respectively) based on the chemical shifts observed, while augmenting aggregation via the amyloidogenic core NAC region (61-95) (Figure 1c). This is readily comprehensible given that the N- and C-terminal ends of αS are hydrophilic and acidic, they could interact electrostatically with highly positively charged TDP-43 PrLD bringing the amyloidogenic NAC region of αS in close proximity to augment aggregation. Unfortunately, similar structural information on TDP-43 PrLD was undecipherable due to rapid aggregation of TDP-43 PrLD and consequent dilution of chemical shifts. Collectively, these data bring forth the synergistic yet selective interactions between broad categories of αS and TDP-43 PrLD species. Yet another significant aspect that highlights the interaction between the two proteins is the ability of αS to modulate LLPS of TDP-43 PrLD and RNA and promote insoluble aggregates. Since these droplets are the main constituents of cytoplasmic SGs, these results demonstrate crucial yet hitherto unseen interactions that may hold significance in neurodegenerative diseases with co-morbidities. Perhaps the most significant of all is the observation that only the hybrid aggregates show far greater cellular toxicity as opposed to the individual aggregates, which suggests that co-morbidities in these pathologies are better defined by the cross-talk between TDP-43 and αS.

**Figure 7.**
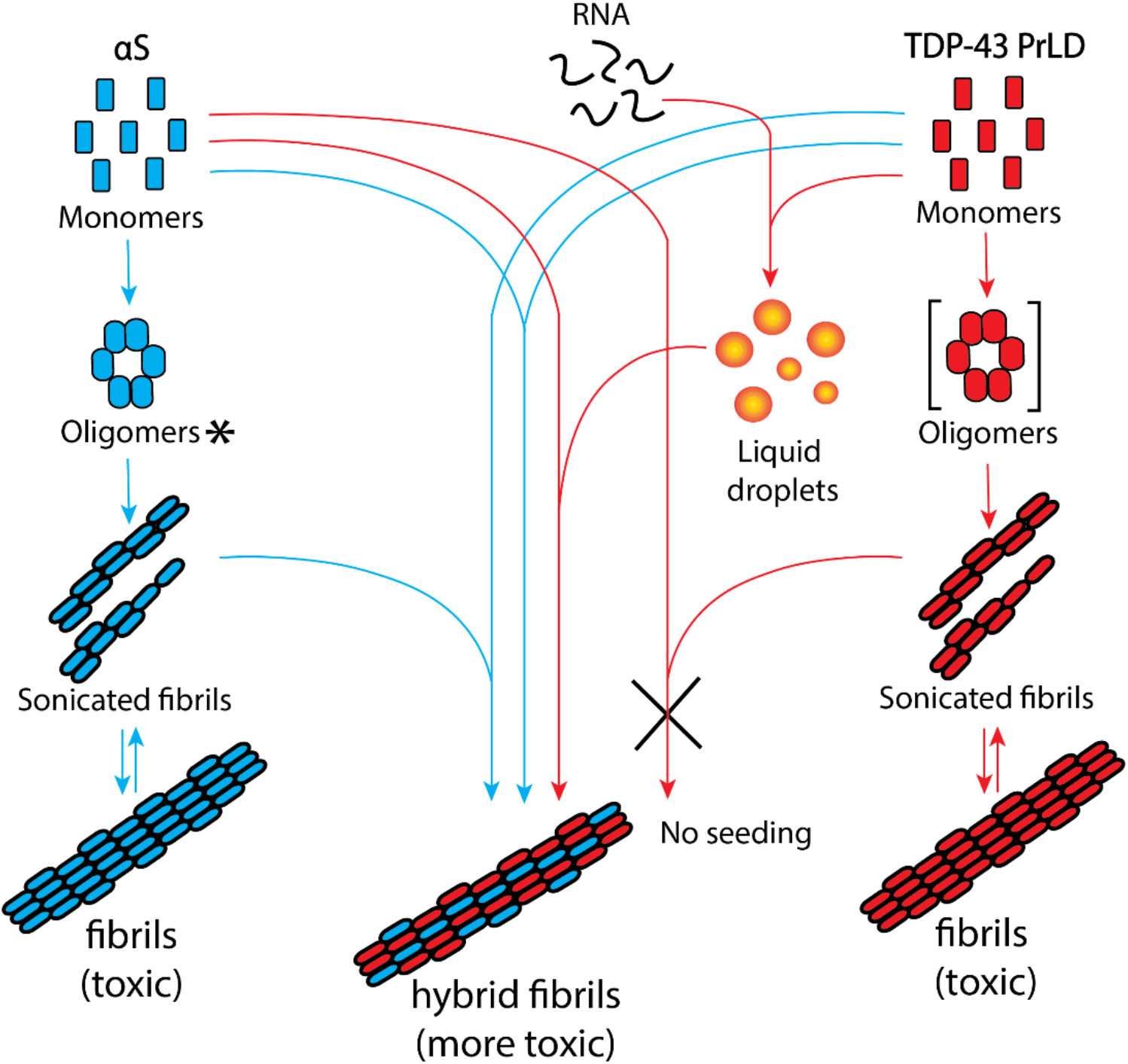
Schematic diagram summarizing the collective results from this work. The square parenthesis indicates theoretical transient oligomers and not used in this study. ‘*’ DOPAL-derived *de facto* intermediate oligomers but shown along the aggregation pathway for simplicity.

It is well-known that aggregates of αS and TDP-43 individually are observed in many neurodegenerative pathologies including AD, PD, Huntington’s disease, FTD, etc. [11, 58-60]. The term ‘synucleinopathies’ has come to define some of the pathologies in which αS aggregates play a causative or a propagative role(s). In the last decade, aggregates of TDP-43 have also been increasingly observed in as many pathologies in which αS aggregates have been observed [15] invoking a compelling argument to categorize some of these maladies as TDP-43 proteinopathies. However, more interesting is the significant overlap between pathologies in which both aggregates of TDP-43 and αS aggregates are observed. Indeed, many reports have indicated the colocalization of TDP-43 and αS aggregates [27, 61] implicating potential interactions between the two proteins to play a role in these pathologies. The data presented here unequivocally demonstrates the interactions between these two proteins which may underlie the clinical and pathological observations, and open doors to deeper investigations in establishing mechanistic links to co-morbidities in neurodegenerative diseases.

## METHODS

### Expression and purification of recombinant proteins

Expression and purification of both unlabeled and ^15^N labeled recombinant TDP-43 PrLD was performed as described previously [40]. Briefly, TDP-43 PrLD fusion construct (Addgene plasmid #98669) was expressed in BL21 Star™ (DE3) cells (Life Technologies) and purified using Ni-NTA affinity chromatography. Concentrated aliquots of protein in storage buffer (20 mM Tris pH 8.0, 500 mM NaCl, and 2M Urea) at −80 DC was thawed on ice and desalted in 20 mM MES buffer pH 6.0 using Zeba Desalting Spin Columns (Thermo) for experimental studies.

Both unlabeled and ^15^N labeled recombinant full-length αS was expressed and purified in Rosetta™ 2(DE3) pLysS (EMD Millipore^®^) cells using IMPACT™ system protocol (New England Biolabs^®^) as described before [62], with few modifications. Briefly, αS detached from chitin beads using dithiothreitol (DTT) was eluted in elution buffer (20 mM Tris, 100 mM NaCl, 1 mM EDTA at pH 8.0). The eluted fraction was filtered using 30 kDa centrifugal filter units (Thermo) at 7000xg for 20 minutes. The flow-through was dialyzed against 4L of nanopure water in a 10 kDa dialysis bag to remove DTT and concentrated using a vacufuge. Concentrated αS was directly subjected to reverse phase HPLC purification by the gradient elution of water and acetonitrile (ACN) each containing 0.1% TFA. HPLC fractions containing pure protein were lyophilized and stored at −80 °C. Lyophilized samples were resuspended in 20 mM MES pH 6.0 and subjected to size exclusion chromatography (SEC) to remove any preformed aggregates and thus obtained pure monomeric protein was used as such for studies. For ^15^N HMQC spectroscopy, TDP-43 PrLD was cleaved by TEV protease, and cross-peaks of αS and TDP-43 PrLD were identified based on previous studies [47, 55, 63].

### Preparation of αS oligomers, αS fibrils and TDP-43 PrLD fibrils

αS oligomers and fibrils were generated similar to previous protocols [64, 65]. Briefly, αS oligomers have been generated by incubating 50 μM αS monomer with 20-fold molar excess of 3,4-Dihydroxyphenylacetaldehyde (DOPAL) at 600 rpm at 37 °C for 24 hours. Oligomers of αS were isolated using superdex-200 size exclusion chromatography (SEC) column in 10 mM Tris pH 8.0. Fibrils of αS were prepared by incubating 5 mg of monomeric protein in presence of 150 mM NaCl at 600 rpm at 37 °C for 7 days. Similarly, TDP-43 PrLD fibrils were generated by incubating 50 μM monomeric TDP-43 in quiescent condition at 37 °C for 7 days. Both fibrils were isolated by centrifuging at 20,000 xg for 20 minutes and stored at −80 °C until use.

### Thioflavin-T (ThT) fluorescence

Aggregation kinetics were monitored using BioTek Synergy H1 microplate reader. Samples containing 10 μM ThT were excited at 452 nm and emission was monitored at 485 nm at 37 °C, and the data obtained was plotted using Origin 8.5.

### SDS-PAGE and immnoblotting

Aliquots of the reactions were subjected to SDS-PAGE and immunoblotting using a monoclonal anti-αS antibody, clone Syn211 (Millipore Sigma). Aliquots of the samples were separately mixed in the 4x Laemmli sample buffer and loaded onto SDS-PAGE Biorad Mini-PROTEAN® 4-20% precast gel. Gels were then transferred on to a 0.45 μM Amersham Protran Premium nitrocellulose membrane (GE Life Sciences) and the blot was boiled in 1X PBS for one minute. Blot was then incubated overnight in the blocking buffer (5% non-fat milk, 0.1% Tween®-20 in 1X PBS), followed by primary antibodies against αS or TDP-43 PrLD and horseradish peroxidase-conjugated anti-mouse/anti-rabbit secondary antibodies. Finally, images were obtained by treating with ECL reagent using GelDoc molecular imager (Bio-Rad).

### MALDI-ToF mass spectrometry

Quantification of αS and TDP-43 PrLD fibrils was performed on a Bruker Datonics Microflex LT/SH MALDI-ToF system. Aliquots of samples from aggregation reactions after 48 hours were centrifuged at 18,000xg for 20 minutes. The supernatant was discarded and the pellet was washed and resuspended in an equal volume of 20 mM MES buffer pH 6.0. Resuspended pellets were then mixed with an equal volume of formic acid to disaggregate the fibrils. The samples were then mixed with Cytochrome C external standard at 1.42 μM final concentration in 1:1 sinnapinic acid matrix and loaded on to an MSP 96 BC MALDI plate (Bruker Datonics). The instrument was calibrated using Bruker Protein Calibration Standard I (Bruker Datonics) and spectra were collected by adjusting the laser intensity at 70%.

### Circular dichroism

Circular dichroism (CD) spectra of 4.25 μM DOPAL-derived αS oligomers in 10 μM Tris buffer pH 8.0 was measured in far UV region (190 to 260 nm) in Jasco J-815 spectrophotometer (Jasco MD) as previously [66].

### Fluorescence Microscopy

Fluorescence microscopic images of the reaction with labeled proteins were obtained using Leica SP8 confocal microscope at 40x magnification in a clear glass bottom 96 well black plates (P96-1.5H-N, Cellvis Inc.). Briefly, protein labeling was carried out by incubating three molar excess of fluorescent dyes, Hilyte 405 or Hilyte 647 (AnaSpec Inc) with the proteins for 12 hours at 4°C. Excess dye was removed using PD SpinTrap™ G-25 (Cytiva Life sciences) columns and the labeled and unlabeled protein samples were mixed in 1:99 ratio and used for the experiments. For thioflavin-S (ThS) staining, aliquots of samples from the reactions were centrifuged at 18,000xg for 20 minutes, and pellets were resuspended in 20 mM MES buffer pH 6.0. ThS was added at a final concentration of 10 μM and incubated for 10 minutes prior to imaging.

### Fluorescence recovery after photobleaching (FRAP)

Hilyte 647 labeled TDP-43 PrLD and RNA reaction with or without Hilyte 405 labeled αS was monitored using FRAP. The samples were photobleached using 100% laser intensity for 10 seconds with 141 iterations and recovery was monitored for 45 seconds. The FRAP kinetic data was plotted as normalized fluorescence with reference to TDP-43 PrLD and RNA control and was fit using Boltzmann function on Origin 8.5.

### NMR spectroscopy

The heteronuclear multiple quantum coherence (HMQC) NMR spectra of ^15^N labeled αS in 20 mM MES, pH 6.0 with 10% D_2_O was acquired after incubation with and without unlabeled TDP-43 PrLD protein. Similarly, ^15^N labeled TDP-43 PrLD NMR spectra in the same buffer was acquired in the presence and absence of unlabeled αS. The data were acquired on a Bruker Advance-III-HD 850 MHz NMR spectrometer equipped with a Bruker TCI cryoprobe at the high-field NMR facility of the University of Alabama, Birmingham as described previously [40].

### Atomic force microscopy

AFM images were obtained following a previously published method [67]. Briefly, mica was cleaved using tape then attached to a magnet. The mica was then treated with 150 μL of 3-aminopropyltriethoxysilane (APTES) solution (500 μL of 3-aminopropyltriethoxysilane in 50 mL of 1 mM acetic acid) for 30 minutes. The APTES solution was then decanted off and the mica substrate was rinsed three times with nanopure H_2_O, dried with N_2,_ and stored for an hour. The 1-4 μM reaction samples were diluted to 100-folds and a volume of 150 μL of the sample solution was deposited onto the mica surface and were allowed to absorb for 30 minutes. The sample solution was then decanted from the mica surface and washed with 150 µL of nanopure H_2_O three times, dried with N_2,_ and stored in a desiccator until imaging. AFM analysis was performed using a Dimension Icon atomic force microscope (Bruker) in PeakForce Tapping mode. AFM scanning was performed using NanoScope 8.15r3sr5 software and the images were analyzed in NanoScope Analysis 1.50 software. Imaging was performed using a sharp silicon nitride cantilever (SNL-C, nominal tip radius of 2 nm; nominal resonance frequency of 56 kHz; nominal spring constant of 0.24 N/m) and a standard probe holder under ambient conditions with 512 × 512 data point resolution. Multiple areas of the mica surface were analyzed, height and phase images were obtained simultaneously, and representative images are reported.

### Cell viability XTT assay

Human neuroblastoma SH-SY5Y cells (ATCC, Manassas, VA) were maintained in a humidified incubator at 37 °C with 5.5% CO_2_ in 1:1 mixture of DMEM and Ham’s F12K medium with 10% FBS and 1% penicillin/streptomycin. Approximately, 15,000 cells were plated in a clear bottom 96 well black plates (Thermo Scientific) 24 hours prior to sample treatment. All the reaction samples were prepared and purified using autoclaved water and buffer to avoid bacterial contamination. Cell medium from wells was replaced with the reaction samples resuspended in complete growth medium and incubated for 24 hours. Cell viability assay was measured using 2,3-bis(2-methoxy-4-nitro-5-sulfophenyl)-5-[(phenylamino)carbonyl]-2H-tetrazolium hydroxide (XTT) assay.

## ACKNOWLEDGEMENTS

The authors would like to thank the following agencies for financial support: National Institute of Aging (1R56AG062292-01) and the National Science Foundation (NSF CBET 1802793) to VR. The authors also thank the National Center for Research Resources (5P20RR01647-11) and the National Institute of General Medical Sciences (8 P20 GM103476-11) from the National Institutes of Health for funding through INBRE for the use of their core facilities.

## DECLARATION OF COMPETING INTEREST

The authors declare that they have no conflicts of interest with the contents of this article.

## SUPPLEMENTARY FIGURES

**Figure S1.**
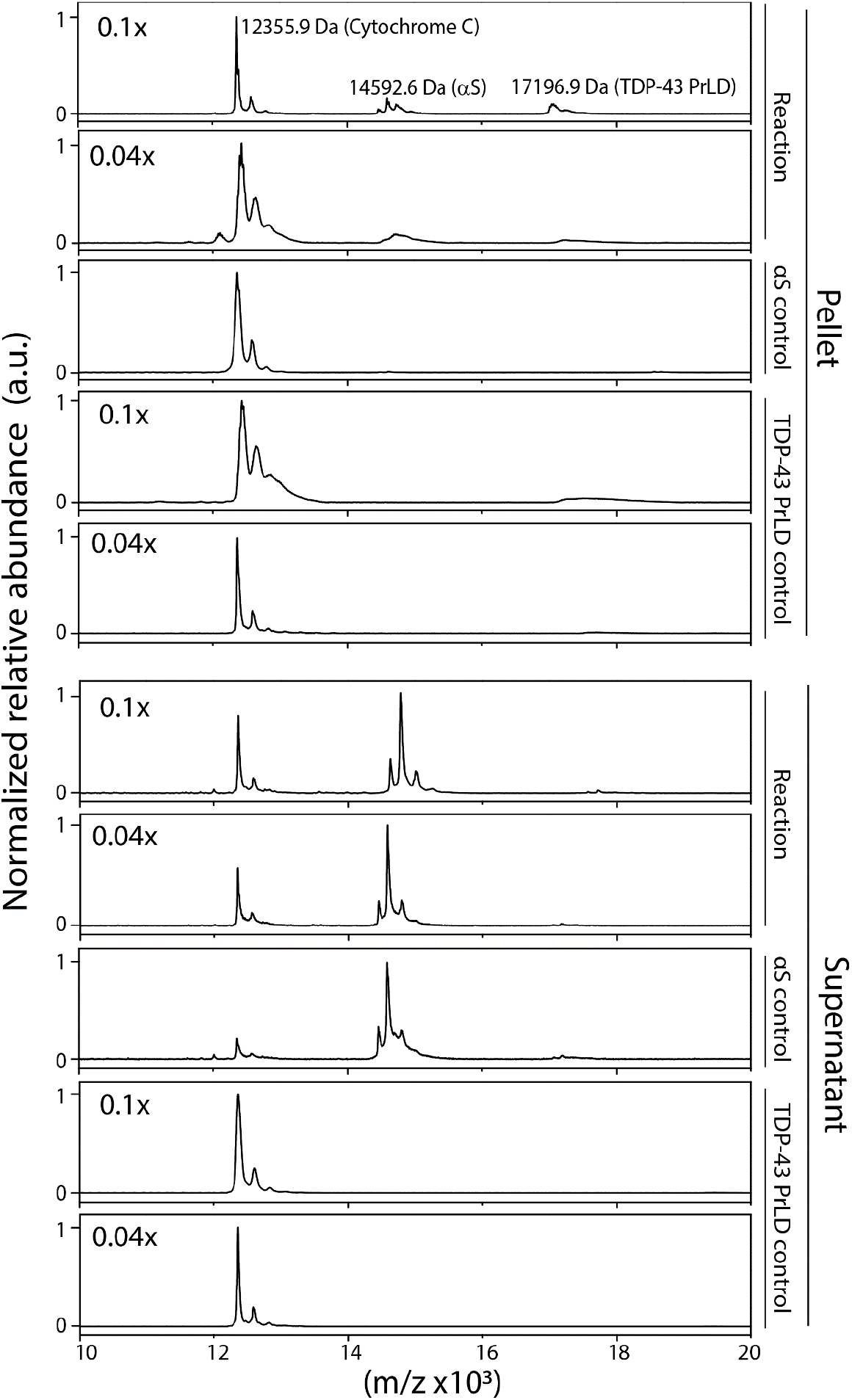
Quantification of αS and TDP-43 PrLD in the supernatant and pellet of reactions in Figure 32 by MALDI ToF. Spectra were obtained after 7248 hours, the reaction samples were centrifuged at 18,000 xg for 20 minutes and analyzed with reference to cytochrome-C as an external standard. (experimental details in materials and methods)

**Figure S2.**
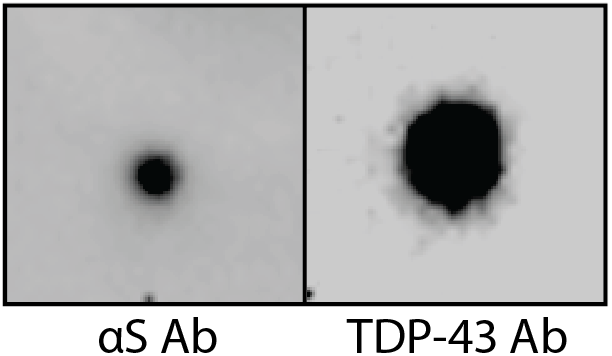
Dot blot analysis of the pellet of reaction containing 20 μM TDP-43 and 2 μM αS in 20 mM MES buffer pH 6.0. Reactions were incubated at 37 °C for 24 hours, centrifuged at 18,000 xg for 20 minutes and analyzed using both αS monoclonal antibody and TDP-43 polyclonal antibody (LifeSpan BioSciences, inc.).

## Notes

### Competing Interest Statement

The authors have declared no competing interest.

